# Explainable machine learning prediction of synergistic drug combinations for precision cancer medicine

**DOI:** 10.1101/331769

**Authors:** Joseph D. Janizek, Safiye Celik, Su-In Lee

## Abstract

Although combination therapy has been a mainstay of cancer treatment for decades, it remains challenging, both to identify novel effective combinations of drugs and to determine the optimal combination for a particular patient’s tumor. While there have been several recent efforts to test drug combinations *in vitro*, examining the immense space of possible combinations is far from being feasible. Thus, it is crucial to develop datadriven techniques to computationally identify the optimal drug combination for a patient. We introduce TreeCombo, an extreme gradient boosted tree-based approach to predict synergy of novel drug combinations, using chemical and physical properties of drugs and gene expression levels of cell lines as features. We find that TreeCombo significantly outperforms three other state-of-theart approaches, including the recently developed DeepSynergy, which uses the same set of features to predict synergy using deep neural networks. Moreover, we found that the predictions from our approach were interpretable, with genes having well-established links to cancer serving as important features for prediction of drug synergy.

## 1. Introduction

Combination drug therapy, which has been utilized in cancer treatment since the 1960s (DeVita & Schein, 1973), is preferred to monotherapy in most cases for a variety of reasons. It has been shown to overcome inherent patient resistance to anti-cancer drugs in cases where monotherapy cannot, and also to prevent the development of acquired drug resistance (Lopez & Banerji, 2017). It also has been shown to lead to a decrease in dose-related toxicities while increasing cancer cell elimination through additive or synergistic effects (Chabner & Thompson, 2018). However, finding new effective combinations of drugs is a complex undertaking since there exists a huge number of possible drug combinations and this number increases each time a new drug is developed. The current strategy for discovering effective drug combinations is largely based on physicians’ experience as they try new combinations in clinic; patient’s molecular data is rarely utilized (Day & Siu, 2016).

While the space of possible drug combinations is too large to be tested exhaustively, there have been recent efforts to measure the efficacy of drug combinations via high-throughput screening (O’Neil et al., 2016; Menden et al., 2018). However, it is unfeasible to exhaustively test the immense space of possible combinations, which clearly motivates the need for a data-driven approach to discovering effective combinations of drugs. The aforementioned datasets from *in vitro* screens enabled development of such approaches, and there have been a variety of prior attempts to use machine learning methods to predict the most synergistic combinations of anti-cancer drugs (Li et al., 2015). A recent study (Preuer et al., 2018) improved predictions by applying deep learning to a large dataset of drug combinations from Merck (O’Neil et al., 2016).

We present TreeCombo, which aims to predict the synergy scores of drug combinations using extreme gradient boosted trees (XGBoost) (Chen & Guestrin, 2016), and explain these predictions using a recent feature attribution method developed for tree models (Lundberg et al., 2018). When applied to data from cancer cell lines (O’Neil et al., 2016), TreeCombo achieves a 10% performance improvement over the best-performing state-of-the-art approach DeepSynergy. Moreover, the genes highly ranked by TreeCombo are highly relevant to known cancer mechanisms. We believe that TreeCombo exhibits a promising potential for personalized medicine (Nature Medicine, 2017) by enabling: (1) identification of effective novel drug combinations for individual patients based on their molecular profiles and (2) advance our understanding of the mechanisms by which drug synergy occur by interpretable drug synergy predictions.

## 2. Methods

### 2.1. Background

XGBoost (Chen & Guestrin, 2016) is a relatively recent machine learning library designed to provide “efficient, flexible and portable” implementations of gradient boosted trees.XGBoost is based on ensembles of classification and regression trees (CARTs), which are obtained by recursively partitioning input data and fitting a real-valued prediction model within each partition. Successive trees are fit on the residuals of the previous trees, and ensemble predictions are obtained by taking the sum of the weighted scores predicted by each tree in the model. XGBoost has been shown to be a powerful prediction model for structured data in various applications (Aibar et al., 2017; Rothschild et al., 2018; de Wiele, 2017).

To interpret predictions of TreeCombo, we used TreeSHAP, an algorithm that calculates fast exact tree solutions for SHAP (SHapley Additive exPlanation) values (Lundberg et al., 2017). These feature attribution values have the advantage of being guaranteed to be the unique solutions that are consistent (i.e., their value never decreases when the true impact of that feature is increased) and locally accurate.

### 2.2. Data

We trained TreeCombo on the high-throughput combination screening data from O’Neil et al. (2016). This data consists of over 22,000 samples, where each sample is one of 583 two-drug combinations tested in 39 cancer cell lines from different tissues of origin. For each sample, cell line viability was measured in response to a four-by-four dosing regimen of a unique 2-drug combination. From these measurements, drug synergy values were calculated according to a Loewe additivity model as described in Preuer et al. (2018) and standardized (i.e., made zero-mean and unit variance). As input features, we used drug physical and chemical features (e.g., molecular connectivity fingerprints, presence or absence of toxicophore structures) and cell line gene expression levels as used by Preuer et al. (2018). Filtering out features with no variance across samples led to 2,431 features per drug and 3,984 features per cell line. Thus, each sample, consisting of a cell line and a 2-drug combination, was described by a total of 8,846 features. Gene expression levels, which were measured using Affymetrix HG-U219 arrays, were accessed from ArrayExpress (http://www.ebi.ac.uk/arrayexpress) with accession number E-MTAB-3610.

### 2.3. Experimental Setup

We compared TreeCombo to: (1) Elastic Net, a regularized linear regression method, (2) Random Forest which uses ensembles of trees like TreeCombo, and (3) DeepSynergy which uses deep neural networks (DNNs). We used scikitlearn (Pedregosa et al., 2011) implementations of Elastic Net and Random Forest. We recreated the DeepSynergy model in Keras (Chollet et al., 2015) with TensorFlow backend, using the architecture described by Preuer et al. (2018).

To ensure that our models generalized to unseen combinations of drugs, we tested TreeCombo and the alternative methods using five-fold cross-validation experiments. To enable comparison of the performance of our model to the performance of DeepSynergy (Preuer et al., 2018), we stratified the data in the same way as that study: for each of the 583 unique combinations of two anti-cancer drugs, we ensured that each combination only appeared in one of the five folds. Then, for each of the five held-out test folds, we trained TreeCombo and the alternative methods using the samples from the remaining four test folds, and predicted the synergy scores for the samples in the test fold.

To determine the best hyperparameters for each of the four models, we tuned the models using a separate validation dataset for each fold. These validation sets each consisted of 25% of the training data that had also been stratified to contain unique drug combinations that were not present in the rest of the training set. For ElasticNet, we tuned *α*, the mixing parameter determining the weights of L1 vs. L2 regularization; for Random Forest, we tuned the number of used trees; for DeepSynergy, we looked at the ten best-performing hyperparameter settings for the DNN as reported in (Preuer et al., 2018). The ten best hyperparameter settings for DeepSynergy had been obtained by an exhaustive tuning over a wide range of possible hyperparameters, including three different schemes for preprocessing features, nine different network architectures, four different learning rates, and two different dropout settings.

We found that TreeCombo was substantially more robust to hyperparameter changes, which allows the model to be tested in different settings much more quickly. For TreeCombo, we tuned our model over several maximum tree depths (4, 6, 8, 10, 12) and learning rates (0.05, 0.10, 0.15). The best performance on the validation set was attained using a maximum tree depth of 6, a learning rate of 0.05, and 1000 estimators, with an early stopping parameter used to prevent overfitting.

We then used TreeSHAP (Lundberg et al., 2018) to calculate feature importance values for each of our predictions in each test fold and retrained models using only the *n* most important features, for varying *n*. We observed that TreeCombo’s performance only slightly decreased even when most of the least important features were dropped. We then performed a literature search for the genes with the highest importance averaged over five folds.

## 3. Results

### 3.1. Prediction Performance

To evaluate the performance of our model, we compared TreeCombo to the following methods for synergy prediction:(1) ElasticNet, a regularized linear regression method, (2) Random Forests, an ensemble machine learning method,and (3) DeepSynergy, a recently published deep approach to the drug synergy prediction problem. pared these approaches by two different evalua sures: (1) mean squared error (MSE) and (2) rank tion of actual synergy scores vs. predicted synerg Prediction quality was averaged across five test f five fold cross-validation experiment (See the details).

Table 1 compares different methods’ performan dicting drug synergy. Averaged across the whole TreeCombo significantly outperformed the three models. Additionally, for each of the five folds left-out test data, TreeCombo outperformed all alt When measured by MSE, TreeCombo’s predic proved by 10% over the next best model, and w sured by rank correlation, TreeCombo’s predic proved by 6% over the next best model.

**Table 1.**
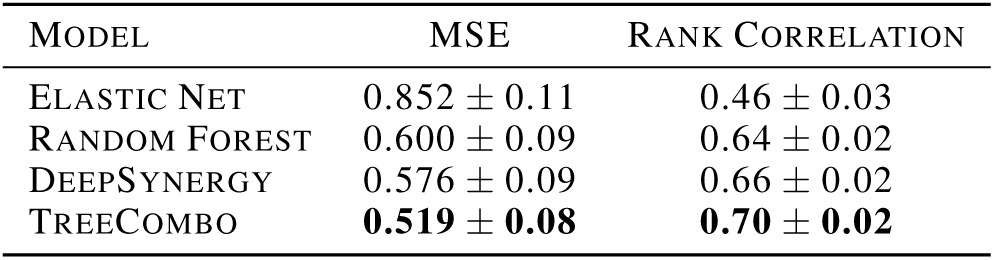
Comparison of methods based on average pre formance across five folds ± one standard deviation across folds

| MODEL | MSE | RANK CORRELATION |
| --- | --- | --- |
| ELASTIC NET | $0.852 \pm 0.11$ | $0.46 \pm 0.03$ |
| RANDOM FOREST | $0.600 \pm 0.09$ | $0.64 \pm 0.02$ |
| DEEPSYNERGY | $0.576 \pm 0.09$ | $0.66 \pm 0.02$ |
| TREECOMBO | **$0.519 \pm 0.08$** | **$0.70 \pm 0.02$** |

To further investigate the quality of predictions TreeCombo, we compared the distributions of predicted synergy scores to the distributions of actual synergy scores by cell lines (Figure 1a,b). While the prediction MSEs varied across cell lines, with some cell lines being predicted more accurately than others, the synergy distributions were captured well and the median MSEs were very similar between the predicted and actual scores across cell lines. To see how well our model predicted the synergy ranking of different combinations of drugs within cell lines, we also plotted the Spearman correlation between TreeCombo predictions and the actual synergy scores by cell line (Figure 1c). The ordering of the cell lines in Figure 1c is the same as in Figure 1a,b, and we observed that the ranking of drug combinations were not predicted more poorly in the cell lines with high MSE, and that the correlations were fairly consistent across all cell lines, predominantly ranging between 0.6 and 0.75.

**Figure 1.**
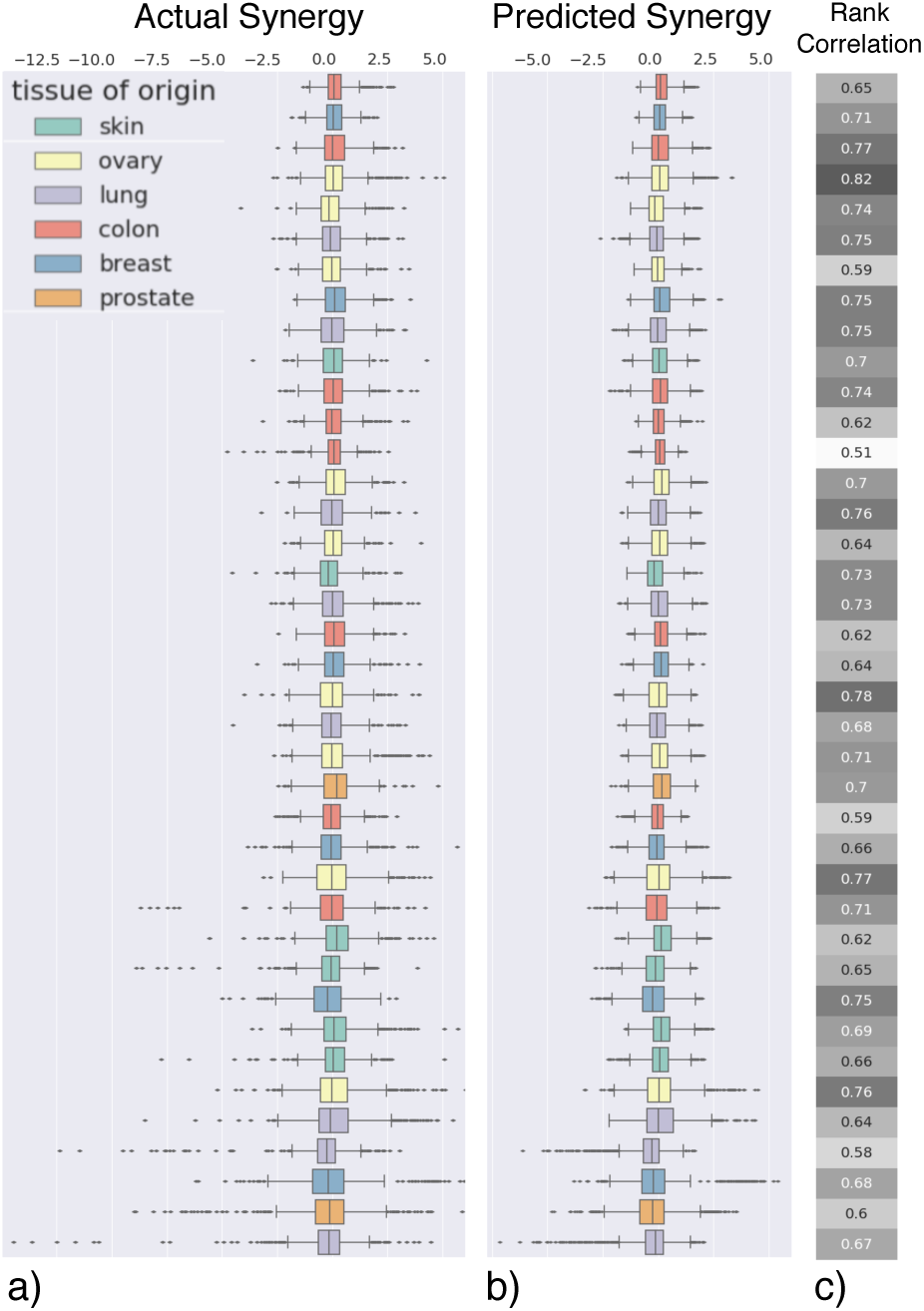
(a, b and c) Box plots of the distribution of the actual (as provided by O’Neil et al. (2016)) vs. TreeCombo-predicted synergy scores for each cell line. Each point represents the measured or predicted synergy score of a unique two-drug combination. Cell lines in both plots are ordered along the y-axis by their MSE as measured in one fold of the held out test data. The rank correlation column shows Spearman’s correlation value between the actual and predicted synergy scores for each cell line.

### 3.2. Feature Selection

One major advantage of using a tree-based method to model our data is the ease of interpretability of our model using the feature attribution method TreeSHAP (Lundberg et al., 2018). TreeSHAP allows for the calculation of fast exact solutions for the unique feature attribution values guaranteed to be consistent and locally accurate. For each of the five models trained for TreeCombo (one for each of the five heldout test folds), we calculated the SHAP values for all of our features. We then selected the most important features for each independent model by selecting the features with the largest average magnitude over all predictions. Using only the top 1,000 or 2,000 features (11% and 22% of all features, respectively), we re-trained the models. We observed that performance is well-preserved using only this small subset of features (Table 2), indicating that the features highly ranked by TreeSHAP were truly important for an accurate prediction. When we used 2,000 most important features selected by TreeSHAP to retrain the models, we observed only a 1.2% increase in mean MSE across five folds, while with 2,000 features at random, we observed a 6.5% increase.

**Table 2.** TreeCombo performance using a subset of features.

| FEATURES USED | MSE | PERFORMANCE DROP |
| --- | --- | --- |
| ALL | 0.519 | — |
| 2000 FROM TREESHAP | 0.525 | 1.16% |
| 2000 BY RANDOM | 0.553 | 6.55% |
| 1000 FROM TREESHAP | 0.528 | 1.73% |
| 1000 BY RANDOM | 0.543 | 4.62% |

### 3.3. Feature Interpretability

#### Most Important Features

While it is important to be able to identify drug combinations that are likely to be synergistic, it is also important to understand *why* our model predicts the synergy of these combinations to be high. Thus, we examined 100 most important features based on their importance identified by TreeSHAP and averaged across all predictions and folds to determine their plausibility as predictors of synergistic anti-cancer effects.

Of the 100 features with highest mean importance across all folds and samples, 83 were drug-based features. These predominantly included structural molecular descriptors like 3D-MoRSE descriptors and the eigenvalues of the drug connectivity matrix. The remaining 17 most important features were expression levels of genes, seven (KLF6, CRIP2, RPS11, CTSH, ONECUT2, SNHG8, and CDH3) of which had been linked to cancer in various studies (Hoffmann et al., 2016; Lo et al., 2011; Cheung et al., 2011; Rauch et al., 2006; Sun et al., 2014). The fact that KLF6, a well-known tumor suppressor, was assigned a large feature importance exhibited a high biological plausibility. KLF6 expression levels have been linked to cancers from many different tissues present in our dataset, including breast cancer (Hatami et al., 2013), colorectal cancer (Reeves et al., 2004), skin cancer (Cai et al., 2014), prostate cancer (Chiam et al.), and lung cancer (Ito et al., 2004).

#### Combination-specific Features

We also examined feature importances at the level of individual drug combinations. For example, sorafenib and erlotinib are used in combination to treat non-small cell lung cancer (Lim et al., 2016). Erlotinib specifically targets the epidermal growth factor receptor, while sorafenib targets the vascular endothelial growth factor (VEGF) receptor. For this combination of drugs, the most important gene expression feature for predicting synergy in our model was epithelial membrane protein 2 (EMP2), a gene whose expression positively regulates VEGF (Gordon et al., 2013). EMP2 expression was not in the 100 most important features when averaged over all combinations, showing the power of a method for which individual prediction-level feature attribution can be applied.

#### Explanation-based Clustering

Finally, we examined whether clustering the genes by their feature importance values (identified by TreeSHAP) across different drug combinations would lead to biologically meaningful groups. Gene expression features with similar importances across drug combinations would be expected to share similar biological functions or pathways which would be targeted by these drug combinations. For each of the five folds of our model, we calculated the mean importance of each gene expression feature across cell lines. We then clustered the genes using *k*-means clustering with *k* = 20 such that each cluster contained around 200 genes. Then we tested for enrichment of particular gene ontology (GO) terms within the clusters using Fisher’s exact test with FDR multiple test correction, using the over-representation test tool (Mi et al., 2017) on http://pantherdb.org/. We found that clustering by SHAP values led to biologically interpretable clusters of gene features. For instance, the first cluster was enriched for genes annotated with the GO terms “programmed cell death” and “apoptotic process” (*q* = 2.55 ×10^−3^, 8.52 ×10^−4^). These make sense as pathways that would be important predictors of drug combination synergy, as they influence cells’ susceptibility to being killed. As expected, the GO terms enriched for the second cluster were distinct from the ones enriched for the first cluster, and included terms like “regulation of innate immune response” and “regulation of protein serine/threonine kinase activity” (*q* = 1.26 *×* 10^−2^, 1.29 *×* 10^−2^).

## 4. Discussion

We present TreeCombo, a powerful XGBoost-based approach that outperforms existing machine learning approaches in predicting synergistic combinations of drugs. Beyond its superiority in terms of prediction accuracy, TreeCombo has several advantages over the alternative methods, specifically over the commonly used DNN-based approaches. Tree-based models are substantially easier to prototype compared to DNNs since they require less hyperparameter tuning or feature preprocessing. Moreover, by using a tree-based model, we could easily incorporate feature importances from TreeSHAP into our model. This allowed us to train an almost equally powerful model using only 11% of the provided data and to make straightforward biological interpretations of our results.

There are various directions to improve and extend TreeCombo. Most importantly, we plan to apply it to drug combination screens from primary *patient* cells. Such a model would be more representative of clinical cases and would increase our model’s potential for precision medicine. We also will explore explanation-based biclustering, where feature importances are clustered by both expression feature and drug combination. Testing for over-represented pathways in these biclusters will help elucidate potentially novel molecular mechanisms of the drug synergy phenotype.

